# Hotspot residues and resistance mutations in the nirmatrelvir-binding site of SARS-CoV-2 main protease: Design, identification, and correlation with globally circulating viral genomes

**DOI:** 10.1101/2022.07.12.499687

**Authors:** Aditya K. Padhi, Timir Tripathi

**Affiliations:** Laboratory for Computational Biology & Biomolecular Design, School of Biochemical Engineering, Indian Institute of Technology (BHU), Varanasi-221005, Uttar Pradesh, India; Molecular and Structural Biophysics Laboratory, Department of Biochemistry, North-Eastern Hill University, Shillong-793022, India; Regional Director’s Office, Indira Gandhi National Open University, Regional Centre Kohima, Kenuozou, Kohima-797001, India

## Abstract

Since the onset of the COVID-19 pandemic, SARS-CoV-2 has acquired numerous variations in its intracellular proteins to quickly adapt, become more infectious, and ultimately develop drug resistance by mutating certain hotspot residues. To keep the emerging variants at bay, including Omicron and subvariants, FDA has approved the antiviral nirmatrelvir for mild-to-moderate and high-risk COVID-19 cases. Like other viruses, SARS-CoV-2 could acquire mutations in its main protease (M^pro^) to adapt and develop resistance against nirmatrelvir. Employing a unique high-throughput protein design technique, the hotspot residues and signatures of adaptation of M^pro^ having the highest probability of mutating and rendering nirmatrelvir ineffective were identified. Our results show that ∼40% of the designed mutations in M^pro^ already exist in the globally circulating SARS-CoV-2 lineages. The work provides a first-hand explanation of the resistance mutations in M^pro^ and is crucial in comprehending viral adaptation, robust antiviral design, and surveillance of evolving M^pro^ variations.

---

Pfizer’s Paxlovid (a combination of nirmatrelvir [PF-07321332] tablets and ritonavir tablets) has been approved as the first oral antiviral for the treatment of COVID-19 by the US Food and Drug Administration (FDA). In December 2021, Paxlovid was granted emergency use authorization (EUA) for adults and pediatric patients. Nirmatrelvir is emerging as a useful drug for treating mild-to-moderate and high-risk COVID-19 cases ^1^. It is a potent inhibitor of the SARS-CoV-2 main protease (M^pro^) with an inhibition constant (*K*_i_) of ∼1 nM ^2^, an EC_50_ value of ∼16 nM ^3^, and an IC_50_ value ranging between 22 and 225 nM ^4^. Using a reversible covalent mechanism, nirmatrelvir, via its cyano group, reacts with the catalytic Cys145 residue of the M^pro^. SARS-CoV-2 is continually evolving and generating new sequence variants globally. While the emergence of new variants is inevitable, predicting their evolution, spread, and impact through intense research may definitely help improve countermeasures ^5-6^. In the current study, we used a unique high-throughput ligand-based interface protein design protocol to identify potential mutational hotspots in the nirmatrelvir-binding site in the SARS-CoV-2 M^pro^. Notably, we correlate our findings with the globally circulating viral genomes and show that our design methodology can correctly predict the hotspot residues and adaptable mutations in the M^pro^.

Structural inspection of the nirmatrelvir-M^pro^ crystal structure revealed 25 interacting residues that were subjected to design (Figures 1A & 1B). Employing the resistance scan module of MOE, the 25 nirmatrelvir-interacting M^pro^ residues were mutated to naturally evolving SNPs (Table 1). This led to 210 single point mutants, of which, the relative binding affinities of the mutations to the native M^pro^ (dAffinity) were computed, where a higher positive dAffinity denoted lower affinity with nirmatrelvir, thereby implying easy tolerance of the mutant to nirmatrelvir. The computed dAffinity of 210 single mutants ranged from -1.75 to 1.91 kcal/mol, of which 94 M^pro^ designed mutants exhibited positive values. These results implied tolerated and adaptable mutants (Figure 2). Further, employing a stringent cut-off of dAffinity >1.0 kcal/mol resulted in the identification of 11 mutants (F140I, G143D, S144C, S144F, S144T, S144Y, M165R, E166A, E166G, E166V, and Q192H), implying they are the potential hotspots and preferred candidates in developing tolerance against nirmatrelvir (Figure 3). This highlighted that residue at positions 144 and 166 are more susceptible to developing tolerance and adaptability during evolution than at other positions.

**Figure 1.**
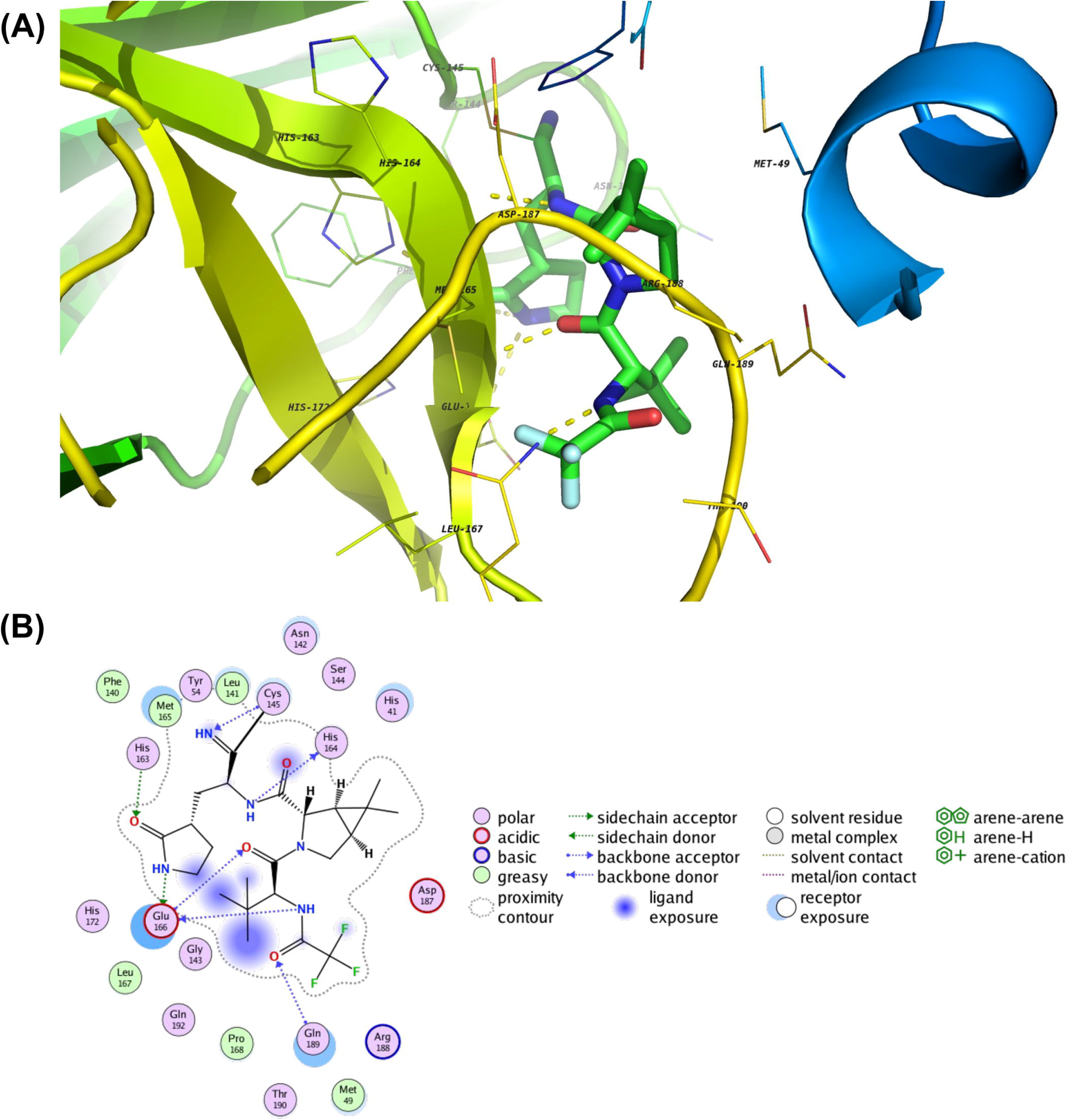
Structure and interactions of the SARS-CoV-2 M^pro^-nirmatrelvir bound complex. (A) The crystal structure of SARS-CoV-2 M^pro^ in complex with nirmatrelvir is shown. The nirmatrelvir is shown as a stick, and the interacting residues of M^pro^ are shown as lines along with hydrogen bond interactions as yellow dashed lines. (B) A 2D representation showing various types of interactions between M^pro^ and nirmatrelvir, where various types of intermolecular interactions are labeled in the legend.

**Table 1.** M^pro^-nirmatrelvir interacting residues that were designed with corresponding SNPs of the native sequence to emulate the mutations that are more likely to happen naturally over the evolution of M^pro^.

| **S. No** | **M<sup>pro</sup>-nirmatrelvir interacting and designed native residues\*** | **Sampled SNPs in designs** |
| --- | --- | --- |
| 1 | Leu27 | RQHIMFPSWV |
| 2 | Cys44 | RGFSWY |
| 3 | Met49 | RILKTV |
| 4 | Pro52 | ARQHLST |
| 5 | Tyr54 | NDCHFS |
| 6 | Phe140 | CILSYV |
| 7 | Leu141 | RQHIMFPSWV |
| 8 | Asn142 | DHIKSTY |
| 9 | Gly143 | ARDCESWV |
| 10 | Ser144 | ACLFPTWY |
| 11 | His163 | RNDQLPY |
| 12 | His164 | RNDQLPY |
| 13 | Met165 | RILKTV |
| 14 | Glu166 | ADQGKV |
| 15 | Leu167 | RQHIMFPSWV |
| 16 | Pro168 | ARQHLST |
| 17 | His172 | RNDQLPY |
| 18 | Phe181 | CILSYV |
| 19 | Val186 | ADEGILMF |
| 20 | Asp187 | ANEGHYV |
| 21 | Arg188 | NCQGHILKMPSTW |
| 22 | Gln189 | REHLKP |
| 23 | Thr190 | ARNIKMPS |
| 24 | Ala191 | DEGPSTV |
| 25 | Gln192 | REHLKP |

\* The catalytic dyad residues His41 and Cys145 of M<sup>pro</sup> were not designed, as they carry out the acylation-deacylation reaction and cleavage of the substrates.

**Figure 2.**
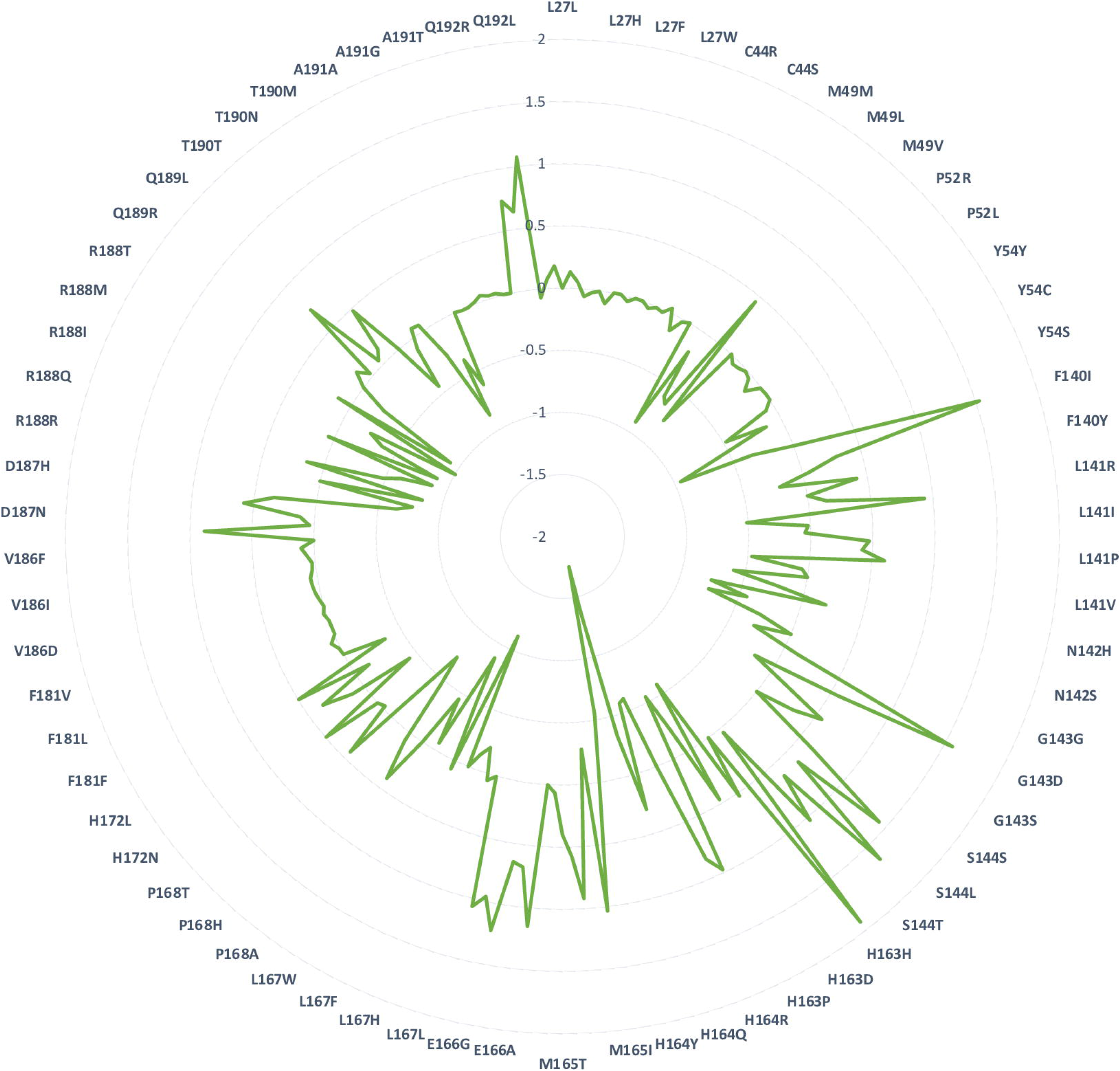
Relative binding affinities (dAffinities) of all the designed mutations in the nirmatrelvir-binding site of M^pro^. Radar plot showing the computed dAffinities for the most plausible tolerated and adaptable single-point mutant designs of M^pro^ bound to nirmatrelvir. The mutated designs are labeled in the radar plot, and the corresponding dAffinity values are shown.

**Figure 3.**
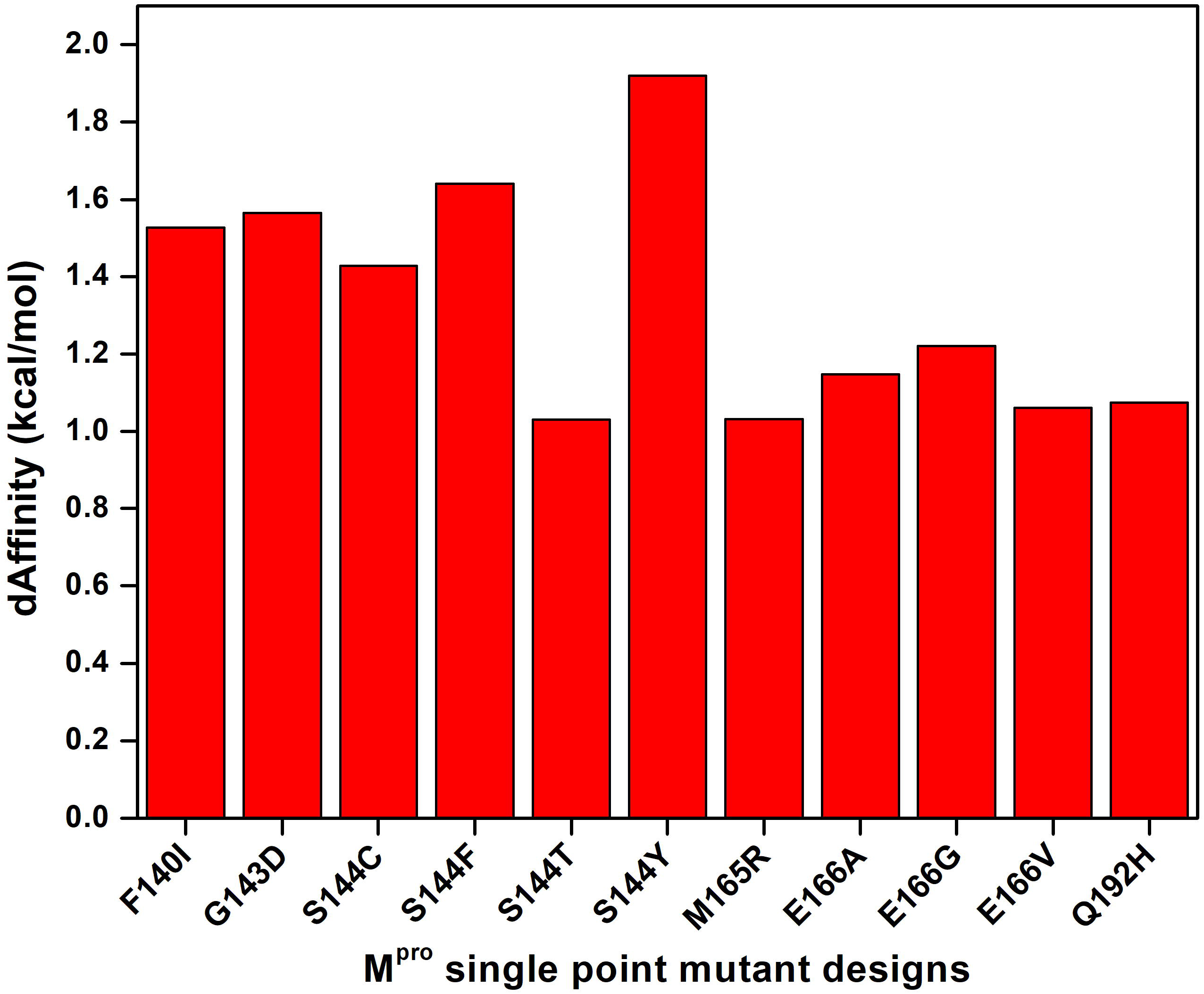
Relative binding affinities (dAffinities) of the most plausible tolerated and resistance-eliciting designs from the M^pro^-nirmatrelvir complex. Bar graph highlighting the dAffinities for the most plausible tolerated and resistance-eliciting mutants of M^pro^ against nirmatrelvir, where designs exhibiting dAffinity > 1.0 kcal/mol suggest a decrease in affinity towards nirmatrelvir and hence their ability to become resistant easily.

Another subsequent analysis focused on evaluating the accuracy of our protein design protocol ^7^ and assessing how closely our designed single-point mutants of M^pro^-nirmatrelvir corroborate with the experimentally determined SARS-CoV-2 sequences from the COVID-19 pandemic. As SARS-CoV-2 is actively mutating due to poor immunity, drug, and vaccine pressures, it resulted in the accumulation of >1000 unique mutations at the M^pro^, thereby continuously being deposited in GISAID and released through CoV-GLUE-Viz. We obtained all the deposited mutations and their frequencies of the nirmatrelvir-bound M^pro^ site and compared them with our designed mutants. The comparison showed that 78 out of 199 mutations were predicted as positively selected and favorable to develop tolerance and adaptability based on our design calculations, thereby achieving ∼40% corroboration with the clinical-sequencing data (Figure 4). Several of the already circulating, high-frequency mutants such as F140C, S144L, P168S, V186F, V186I, T190I, and A191V are already found to be having higher dAffinity values in our design computations. This indicates that these mutants are already tolerant and adaptive against nirmatrelvir, even without significant drug or selection pressure (Figures 3 and 4). Although it is vital to note that the dAffinities values cannot directly be correlated with observed frequencies, some of the designed mutants with dAffinities >1.0 kcal/mol, such as S144C, are already among the high-frequency mutants (Figures 3 and 4). Notably, while this manuscript was being prepared, a preprint article experimentally reported 11 mutants at the drug-binding site of M^pro^ that are resistant to nirmatrelvir (*K*_i_ > 10-fold increase) ^8^. This offered us an opportunity to critically assess the designed M^pro^ mutants with that of the experimentally reported resistant mutants. Upon examination, it was found that several of the designed mutants, such as S144A, S144F, S144Y, M165T, E166Q, and H172Q, were already found to be resistant and naturally occurring, therefore corroborating our findings and reflecting the accuracy of our design methodology.

**Figure 4.**
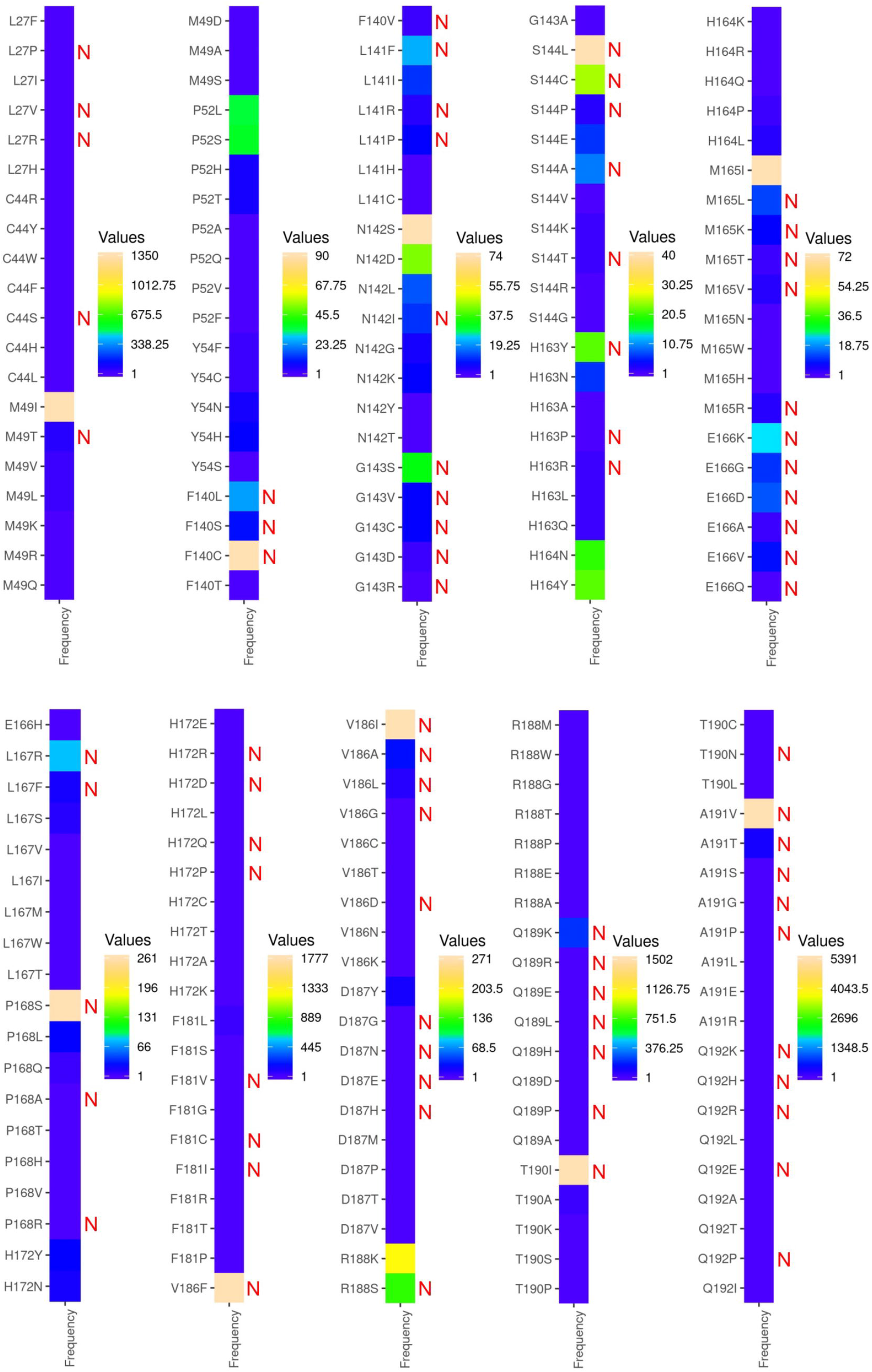
Heat maps of the nirmatrelvir-bound M^pro^ mutations with their frequencies of occurrence retrieved from GISAID-enabled CoV-GLUE-Viz databases. Mutations at the nirmatrelvir-binding site of M^pro^ obtained from the GISAID-enabled CoV-GLUE-Viz databases are shown. Frequencies of the mutations in SARS-CoV-2 sequences from the COVID-19 pandemic ranged from lower to higher numbers and from blue to orange colors, respectively. The designed mutants that potentially developed tolerance and adaptation towards nirmatrelvir are denoted as ‘N’ in red. It was found that out of 199 mutations, 78 mutations were predicted as positively selected and tolerant in the design computations, therefore attaining ∼40% correlation, already exist in the SARS-CoV-2 sequences and are currently circulating.

A further inspection of the high-frequency mutants F140C, S144L, P168S, V186F, V186I, T190I, and A191V with respect to the lineages revealed that these mutants pertain to some of the globally distributed lineages that are found in nearly all the COVID-19 prevalent countries. More specifically, the F140C mutant is detected in six, S144L in 23, P168S in 69, V186F in 102, V186I in 41, T190I in 126, and A191V in 285 different lineages (a total of 652 lineages), out of which 421 lineages (64.5%) were found to be common across all the seven high-frequency mutants (Supplementary Table 1). This suggests that the SARS-CoV-2 strains with certain residue-specific mutations of M^pro^ are already circulating in humans, even without significant nirmatrelvir pressure. Notably, this finding validated our design protocol and highlighted that as the virus experiences more diverse pressure and sequencing data keeps piling up, several of our designed mutations with high dAffinity values may appear extremely valuable in surveillance and predicting drug tolerance and adaptability.

Finally, the intermolecular interactions between nirmatrelvir and M^pro^ in the high-affinity and low-affinity designs were visualized and obtained. While the high-affinity design had 431 interactions, the low-affinity design had only 407 interactions. The proximal and van der Waals interactions were crucial in driving the affinity between nirmatrelvir and M^pro^. The interaction diagrams between the high-affinity and low-affinity design demonstrated that losing a critical hydrogen bond between nirmatrelvir and Cys145 reduces the affinity and develops tolerance and adaptability (Supplementary Figure 1).

Nonetheless, we recognize certain limitations of the study, firstly, the requirement of the experimental and functional validation of the nirmatrelvir-tolerated designs. Secondly, it is also reported that mutations distal from the catalytic or active site (e.g., HIV-1 protease) can also cause drug tolerance, which is not addressed in this work. Finally, despite our recent results identified critical residues in the RNA-dependent RNA polymerase (RdRp) that could render SARS-CoV-2 resistant to remdesivir, molnupiravir, and favipiravir ^9-10^, a more systematic study to develop a unifying methodology is in progress to address some of these challenges.

A recent phase 2–3 double-blind, randomized, controlled trial showed that the administration of Paxlovid to symptomatic COVID-19 patients resulted in an 89% lower risk of disease progression to severe conditions and quickly reduced viral load without evident safety issues ^11-12^. However, the rapid and regular emergence of new SARS-CoV-2 variants highlights the fatal nature of the virus to mutate and illustrates that the COVID-19 pandemic will stay in the foreseeable future. Continuous surveillance and prediction of residues prone to mutation that can render therapies ineffective can emerge as a crucial repository for managing the spread of emerging mutants and controlling the pandemic. Currently, > 3 million SARS-CoV-2 genome sequences have been submitted to the GISAID website. The use of high-throughput technologies can considerably extend the insights gathered during drug development, improve drug efficacy and safety, predict the challenging multiple variants, and strengthen the countermeasures. Therefore, one may either wait for the new tolerant and adaptable variants to develop and then battle it or anticipate it coming and expand the treatment arsenals in advance. With the currently available data and the emergence of new SARS-CoV-2 variants, the scientific community should be prepared to tackle the potential nirmatrelvir tolerant variants and intensify the development of additional interventions.

## Methods

The crystal structure of SARS-CoV-2 M^pro^ in complex with nirmatrelvir (PDB ID: 7TLL) was used to obtain the M^pro^’s interacting residues with nirmatrelvir ^2^ and then prepared using Molecular Operating Environment (MOE), v2022.03. Subsequently, a high-throughput protein design involving the resistance scan methodology of MOE was employed to identify single point mutations of the nirmatrelvir-binding site in M^pro^ that exhibit the highest potential for adaptability and drug tolerance ^7, 13^. To sample the mutations that are not lethal to the virus and more likely to evolve naturally, the nirmatrelvir-binding sites were mutated to single nucleotide polymorphisms (SNPs) of the native sequence only. From the nirmatrelvir-bound M^pro^ complex, 25 drug-binding residues of M^pro^ were designed, except the catalytic dyad (His41 and Cys145) because of their role in substrate cleave and processing ^14-15^. For exhaustive sampling and designed site flexibility, the ensemble protein design protocol with the rotamer explorer option was utilized, followed by the root mean square deviation (RMSD) threshold of 0.5 Å. Other important parameters, including the “conformation limit”, “fix residues farther than,” and “energy window,” were set to 25 K, 4.5 Å, and 10 kcal/mol, respectively. A total of 210 designed mutations were generated, and the relative binding affinity of the mutation to the native M^pro^ (represented by dAffinity, which is the Boltzmann average of the relative affinities of the ensemble) was computed for the designed M^pro^-nirmatrelvir complexes. The Arpeggio webserver was used to analyze the intermolecular interactions between nirmatrelvir and M^pro^ in the high-affinity and low-affinity designs of the M^pro^-nirmatrelvir complex ^16^. Finally, to validate the protein design protocol and correlate the designed M^pro^ mutations with available experimental data, the designed M^pro^ residues that bind to nirmatrelvir were compared with globally circulating SARS-CoV-2 sequences deposited in the Global Initiative on Sharing Avian Influenza Database (GISAID) enabled CoV-GLUE-Viz database, after which their frequencies of occurrence were obtained to demonstrate design accuracy ^17-18^.

## Author Contributions

AKP carried out all the design experiments, data generation, and analysis. AKP and TT conceived the study, participated in its design and coordination, and drafted the manuscript. Both authors read and approved the final manuscript.

## Declaration of competing interest

The authors declare no competing interests.

## Acknowledgments

The authors thank Dr. Kam Y.J. Zhang (Laboratory for Structural Bioinformatics, RIKEN, Yokohama) for his support and valuable suggestions for improving the manuscript. The authors acknowledge RIKEN ACCC for the Hokusai supercomputing resources.

## Notes

### Competing Interest Statement

The authors have declared no competing interest.

## References

1. Owen, D. R.; Allerton, C. M. N.; Anderson, A. S.; Aschenbrenner, L.; Avery, M.; Berritt, S.; Boras, B.; Cardin, R. D.; Carlo, A.; Coffman, K. J.; Dantonio, A.; Di, L.; Eng, H.; Ferre, R.; Gajiwala, K. S.; Gibson, S. A.; Greasley, S. E.; Hurst, B. L.; Kadar, E. P.; Kalgutkar, A. S.; Lee, J. C.; Lee, J.; Liu, W.; Mason, S. W.; Noell, S.; Novak, J. J.; Obach, R. S.; Ogilvie, K.; Patel, N. C.; Pettersson, M.; Rai, D. K.; Reese, M. R.; Sammons, M. F.; Sathish, J. G.; Singh, R. S. P.; Steppan, C. M.; Stewart, A. E.; Tuttle, J. B.; Updyke, L.; Verhoest, P. R.; Wei, L.; Yang, Q.; Zhu, Y., An oral SARS-CoV-2 M(pro) inhibitor clinical candidate for the treatment of COVID-19. Science 2021, 374 (6575), 1586–1593. DOI: 10.1126/science.abl4784.

2. Greasley, S. E.; Noell, S.; Plotnikova, O.; Ferre, R.; Liu, W.; Bolanos, B.; Fennell, K.; Nicki, J.; Craig, T.; Zhu, Y.; Stewart, A. E.; Steppan, C. M., Structural basis for Nirmatrelvir in vitro efficacy against the Omicron variant of SARS-CoV-2. bioRxiv 2022, 2022.01.17.476556. DOI: 10.1101/2022.01.17.476556.

3. Rai, D. K.; Yurgelonis, I.; McMonagle, P.; Rothan, H. A.; Hao, L.; Gribenko, A.; Titova, E.; Kreiswirth, B.; White, K. M.; Zhu, Y.; Anderson, A. S.; Cardin, R. D., Nirmatrelvir, an orally active Mpro inhibitor, is a potent inhibitor of SARS-CoV-2 Variants of Concern. bioRxiv 2022, 2022.01.17.476644. DOI: 10.1101/2022.01.17.476644.

4. Rosales, R.; McGovern, B. L.; Rodriguez, M. L.; Rai, D. K.; Cardin, R. D.; Anderson, A. S.; group, P. S. P. s.; Sordillo, E. M.; van Bakel, H.; Simon, V.; García-Sastre, A.; White, K. M., Nirmatrelvir, Molnupiravir, and Remdesivir maintain potent *in vitro* activity against the SARS-CoV-2 Omicron variant. bioRxiv 2022, 2022.01.17.476685. DOI: 10.1101/2022.01.17.476685.

5. Padhi, A. K.; Tripathi, T., Can SARS-CoV-2 accumulate mutations in the S-protein to increase pathogenicity? ACS Pharmacol Transl Sci 2020, 3 (5), 1023–1026. DOI: 10.1021/acsptsci.0c00113.

6. Padhi, A. K.; Tripathi, T., High-throughput design of symmetrical dimeric SARS-CoV-2 main protease: structural and physical insights into hotspots for adaptation and therapeutics. Phys Chem Chem Phys 2022, 24 (16), 9141–9145. DOI: 10.1039/d2cp00171c.

7. Padhi, A. K.; Tripathi, T., A comprehensive protein design protocol to identify resistance mutations and signatures of adaptation in pathogens. Brief Funct Genomics 2022.

8. Hu, Y.; Lewandowski, E. M.; Tan, H.; Morgan, R. T.; Zhang, X.; Jacobs, L. M. C.; Butler, S. G.; Mongora, M. V.; Choy, J.; Chen, Y.; Wang, J., Naturally occurring mutations of SARS-CoV-2 main protease confer drug resistance to nirmatrelvir. bioRxiv 2022, 2022.06.28.497978. DOI: 10.1101/2022.06.28.497978.

9. Padhi, A. K.; Dandapat, J.; Saudagar, P.; Uversky, V. N.; Tripathi, T., Interface-based design of the favipiravir-binding site in SARS-CoV-2 RNA-dependent RNA polymerase reveals mutations conferring resistance to chain termination. FEBS Lett 2021, 595 (18), 2366–2382. DOI: https://doi.org/10.1002/1873-3468.14182.

10. Padhi, A. K.; Shukla, R.; Saudagar, P.; Tripathi, T., High-throughput rational design of the remdesivir binding site in the RdRp of SARS-CoV-2: implications for potential resistance. iScience 2021, 24 (1), 101992. DOI: 10.1016/j.isci.2020.101992.

11. Hammond, J.; Leister-Tebbe, H.; Gardner, A.; Abreu, P.; Bao, W.; Wisemandle, W.; Baniecki, M.; Hendrick, V. M.; Damle, B.; Simón-Campos, A.; Pypstra, R.; Rusnak, J. M., Oral Nirmatrelvir for High-Risk, Nonhospitalized Adults with Covid-19. New England Journal of Medicine 2022. DOI: 10.1056/NEJMoa2118542.

12. Ullrich, S.; Ekanayake, K. B.; Otting, G.; Nitsche, C., Main protease mutants of SARS-CoV-2 variants remain susceptible to nirmatrelvir. Bioorg Med Chem Lett 2022, 62, 128629. DOI: 10.1016/j.bmcl.2022.128629.

13. Vilar, S.; Cozza, G.; Moro, S., Medicinal chemistry and the molecular operating environment (MOE): application of QSAR and molecular docking to drug discovery. Curr Top Med Chem 2008, 8 (18), 1555–72. DOI: 10.2174/156802608786786624.

14. Hilgenfeld, R., From SARS to MERS: crystallographic studies on coronaviral proteases enable antiviral drug design. Febs j 2014, 281 (18), 4085–96. DOI: 10.1111/febs.12936.

15. Ramos-Guzmán, C. A.; Ruiz-Pernía, J. J.; Tuñón, I., Unraveling the SARS-CoV-2 Main Protease Mechanism Using Multiscale Methods. ACS Catal 2020, 10, 12544–12554. DOI: 10.1021/acscatal.0c03420.

16. Jubb, H. C.; Higueruelo, A. P.; Ochoa-Montaño, B.; Pitt, W. R.; Ascher, D. B.; Blundell, T. L., Arpeggio: A Web Server for Calculating and Visualising Interatomic Interactions in Protein Structures. J Mol Biol 2017, 429 (3), 365–371. DOI: 10.1016/j.jmb.2016.12.004.

17. Elbe, S.; Buckland-Merrett, G., Data, disease and diplomacy: GISAID’s innovative contribution to global health. Glob Chall 2017, 1 (1), 33–46. DOI: 10.1002/gch2.1018.

18. Singer, J.; Gifford, R.; Cotten, M.; Robertson, D., CoV-GLUE: A Web Application for Tracking SARS-CoV-2 Genomic Variation. Preprints 2020, 2020060225. DOI: 10.20944/preprints202006.0225.v1.

