## Supplementary Table 1, Supplementary Figure 1 for "Hotspot residues and resistance mutations in the nirmatrelvir-binding site of SARS-CoV-2 main protease: Design, identification, and correlation with globally circulating viral genomes": Nirmatrelvir-Supplementary-bioRxiv.pdf

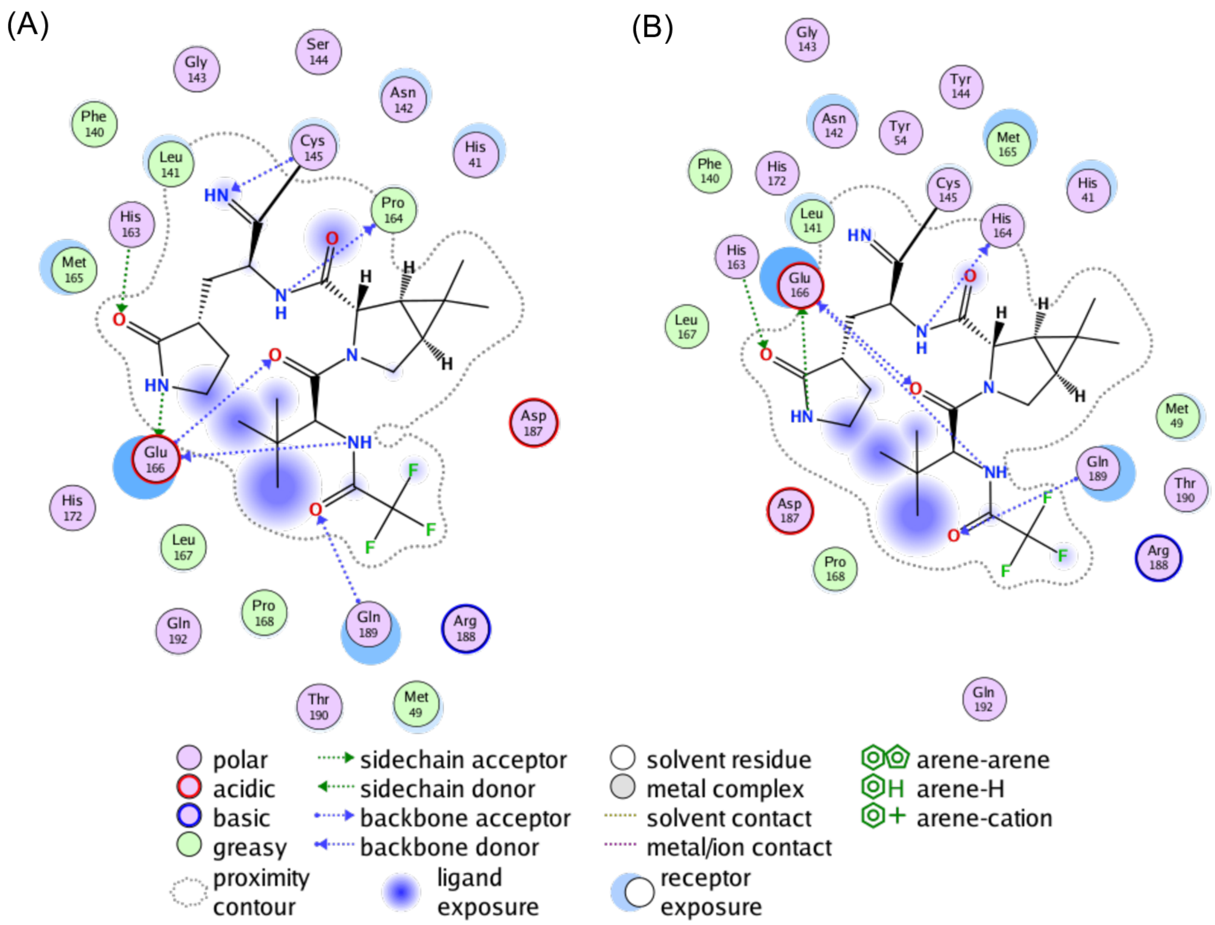


**Supplementary Figure 1. Intermolecular interactions and ligand interaction diagrams between M^pro^-nirmatrelvir for the designs.** 2D ligand interactions between M^pro^ and nirmatrelvir for the (A) high-affinity and (B) low-affinity design. Various types of intermolecular interactions are labeled in the legend.

**Supplementary Table 1.** List of all 652 lineages reported in the GISAID database, where the 7 high-frequency mutants of $M^{pro}$ (F140C, S144L, P168S, V186F, V186I, T190I, and A191V) are found. Among 652 lineages, the 421 commonly occurring lineages across the 7 mutants are highlighted in red cells.

| F140C | S144L | P168S | V186F | V186I | T190I | A191V |
|---|---|---|---|---|---|---|
| AY.9.2 | B.1.1.7 | AY.4 | P.1.13 | AY.23 | AY.4 | B.1.177.20 |
| B.1.1.7 | B.1 | B.1.2 | B.1.427 | AY.39 | B.1.2 | AY.4 |
| B.1.617.1 | AY.25 | B.1 | B.1.1.7 | AY.25 | B.1.1.7 | B.1.1.7 |
| AY.118 | B.1.1 | B.1.160 | AY.4 | AY.120 | B.1.526 | B.1.2 |
| AY.119 | B.1.617.2 | B.1.617.2 | AY.43 | AY.4 | AY.24 | B.1.617.2 |
| AY.75 | P.1.1 | B.1.1 | B.1.240.1 | AY.4.2 | B.1.1.207 | AY.25 |
| | A | AY.4.2 | B.1.617.2 | B.1.1.519 | B.1.617.2 | AY.103 |
| | AY.103 | B.1.1.7 | AY.3 | AY.43 | AY.43 | B.1 |
| | AY.119 | AY.43 | P.1 | AY.44 | AY.11 | B.1.206 |
| | AY.39 | B.1.177 | B.1.438.1 | AY.103 | B.1.575 | B.1.177 |
| | AY.4 | AY.44 | B.1 | AY.6 | AY.25 | AY.43 |
| | AY.9.2 | B.1.177.56 | AY.9.2 | AY.75 | B.1 | AY.44 |
| | B.1.1.317 | B.1.243 | AY.44 | B.1.617.2 | AY.26 | B.1.1.519 |
| | B.1.258 | AY.25 | AY.3.1 | AY.39.1 | AY.3 | B.1.1 |
| | B.1.324 | AY.3 | B.1.160 | AY.74 | B.1.240 | B.1.429 |
| | B.1.336 | AY.39 | AY.102 | AY.45 | A.1 | AY.3 |
| | B.1.351 | B.1.311 | N.5 | B.1.1.7 | AY.44 | AY.42 |
| | B.1.437 | AY.103 | B.1.177.17 | B.1.36.1 | B.1.1.369 | AY.4.3 |
| | B.1.595 | B.1.1.174 | B.1.221 | AY.26 | B.1.243 | AY.5 |
| | B.1.596 | B.1.258.11 | AY.103 | B.1.177 | B.1.1 | B.1.36 |
| | B.31 | AY.50 | B.1.177 | B.1.2 | AY.29 | B.1.1.93 |
| | B.6.6 | B.1.1.214 | B.1.1 | AY.1 | AY.20 | B.1.1.328 |
| | D.2 | B.1.177.54 | B.1.36.29 | AY.42 | B.1.1.279 | B.1.160 |
| | | B.1.429 | AY.45 | B.1 | B.1.177.7 | B.1.351 |
| | | B.1.497 | AY.98 | B.1.1 | AY.103 | B.1.177.21 |
| | | A.2.2 | B.1.1.50 | B.1.562 | AY.47 | AY.85 |
| | | AY.101 | B.1.149 | P.1 | B.1.177 | AY.39 |
| | | AY.102 | B.4 | AY.100 | AY.39 | B.1.526 |
| | | AY.117 | AY.120 | AY.121 | AY.100 | B.1.561 |
| | | AY.20 | AY.20 | AY.47 | B | AY.4.2 |
| | | AY.33 | B.1.170 | AY.68 | B.1.427 | AY.102 |
| | | AY.4.5 | B.1.177.81 | B.1.1.397 | B.1.621 | B.1.1.41 |
| | | AY.46.4 | B.1.2 | B.1.160 | AY.119 | AY.20 |
| | | AY.46.5 | B.1.351.3 | B.1.258 | AY.23 | AY.6 |
| | | AY.47 | B.1.526 | B.1.36 | B.1.429 | B.1.160.9 |

| | | | | | | |
|---|---|---|---|---|---|---|
| | | AY.54 | P.1.1 | B.1.577 | P.1 | B.1.223 |
| | | AY.99.2 | P.2 | B.1.609 | AH.2 | B.1.617.1 |
| | | AZ.1 | AE.8 | B.1.617.1 | B.1.239 | AY.26 |
| | | B | AY.25 | C.31 | AY.118 | AY.29 |
| | | B.1.1.1 | AY.29 | N.3 | AY.120 | AY.39.1 |
| | | B.1.1.316 | AY.36 | P.1.16 | AY.98 | AY.47 |
| | | B.1.1.317 | AY.39 | | B.1.126 | AY.98 |
| | | B.1.1.369 | AY.98.1 | | B.1.177.81 | AY.9 |
| | | B.1.1.39 | B.1.1.216 | | C.23 | AY.100 |
| | | B.1.1.413 | B.1.1.33 | | C.37 | B.1.1.28 |
| | | B.1.1.519 | B.1.177.41 | | AY.102 | B.1.258.17 |
| | | B.1.1.61 | B.1.429 | | AY.120.2 | B.1.596 |
| | | B.1.131 | B.1.470 | | AY.46.6 | R.1 |
| | | B.1.160.16 | B.1.597 | | AY.5 | D.2 |
| | | B.1.177.44 | B.1.621 | | B.1.1.284 | B.1.177.75 |
| | | B.1.177.74 | A | | B.1.351 | P.1 |
| | | B.1.177.79 | AY.101 | | B.1.366 | AY.106 |
| | | B.1.177.87 | AY.116.1 | | Q.2 | AY.46 |
| | | B.1.221.4 | AY.117 | | AY.106 | AY.75 |
| | | B.1.260 | AY.119 | | AY.29.1 | B.1.258 |
| | | B.1.338 | AY.2 | | AY.4.2 | B.1.1.216 |
| | | B.1.351 | AY.23 | | AY.5.4 | B.1.234 |
| | | B.1.396 | AY.24 | | AY.57 | B.1.177.7 |
| | | B.1.399 | AY.26 | | AY.61 | B.1.221 |
| | | B.1.526 | AY.33 | | AY.7.1 | B.1.243 |
| | | B.1.567 | AY.4.2 | | B.1.1.189 | B.1.258.16 |
| | | B.1.575.1 | AY.5.4 | | B.1.1.214 | AY.7.1 |
| | | B.1.576 | AY.59 | | B.1.1.306 | AY.9.2 |
| | | B.1.577 | AY.61 | | B.1.1.519 | AY.96 |
| | | B.1.588 | AY.68 | | B.1.160 | P.2 |
| | | B.1.595 | AY.71 | | B.1.177.60 | B.1.1.434 |
| | | B.1.596 | B.1.1.163 | | B.1.36.26 | B.1.428 |
| | | B.1.621 | B.1.1.207 | | B.1.568 | AY.119 |
| | | B.1.9.5 | B.1.1.214 | | B.1.634 | AY.99.2 |
| | | | B.1.1.222 | | B.29 | B.1.1.214 |
| | | | B.1.1.284 | | Q.7 | B.1.240 |
| | | | B.1.1.294 | | A.27 | AY.120 |
| | | | B.1.1.316 | | A.3 | AY.45 |
| | | | B.1.1.37 | | AD.2 | B.1.1.284 |
| | | | B.1.1.375 | | AY.121 | B.1.177.86 |
| | | | B.1.1.416 | | AY.38 | B.1.466.2 |
| | | | B.1.1.519 | | AY.39.1 | AY.23 |

| | | | | | | |
|---|---|---|---|---|---|---|
| | | | B.1.110.3 | | AY.39.2 | AY.33 |
| | | | B.1.177.12 | | AY.42 | AY.41 |
| | | | B.1.177.15 | | AY.45 | B.1.1.232 |
| | | | B.1.177.21 | | AY.65 | B.1.427 |
| | | | B.1.177.47 | | AY.74 | C.36 |
| | | | B.1.177.82 | | AY.9 | N.8 |
| | | | B.1.221.3 | | AY.90 | AY.46.6 |
| | | | B.1.234 | | AY.98.1 | AY.9.1 |
| | | | B.1.236 | | B.1.1.1 | B.1.1.33 |
| | | | B.1.258 | | B.1.1.141 | B.1.396 |
| | | | B.1.324 | | B.1.1.222 | B.1.609 |
| | | | B.1.351 | | B.1.1.226 | B.39 |
| | | | B.1.36 | | B.1.1.239 | AY.114 |
| | | | B.1.367 | | B.1.1.262 | AY.30 |
| | | | B.1.466.2 | | B.1.1.307 | B.1.232 |
| | | | B.1.551 | | B.1.1.333 | B.1.497 |
| | | | B.1.561 | | B.1.1.351 | B.1.525 |
| | | | B.1.564 | | B.1.1.37 | B.1.532 |
| | | | B.1.595 | | B.1.1.448 | B.1.588 |
| | | | B.1.617.1 | | B.1.1.456 | B.1.637 |
| | | | K.2 | | B.1.1.486 | AY.118 |
| | | | P.1.12 | | B.1.1.518 | AY.36 |
| | | | P.1.17 | | B.1.13 | AY.37 |
| | | | P.1.7 | | B.1.160.16 | AY.46.4 |
| | | | Q.8 | | B.1.177.4 | AY.68 |
| | | | | | B.1.177.57 | AY.84 |
| | | | | | B.1.177.86 | B |
| | | | | | B.1.214.3 | B.1.1.189 |
| | | | | | B.1.234 | B.1.1.222 |
| | | | | | B.1.258.17 | B.1.1.25 |
| | | | | | B.1.258.3 | B.1.1.50 |
| | | | | | B.1.311 | B.1.1.63 |
| | | | | | B.1.351.2 | B.1.22 |
| | | | | | B.1.351.3 | B.1.346 |
| | | | | | B.1.361 | B.1.36.29 |
| | | | | | B.1.367 | B.1.366 |
| | | | | | B.1.400 | B.1.37 |
| | | | | | B.1.408 | B.1.404 |
| | | | | | B.1.564 | B.1.509 |
| | | | | | B.1.574 | B.1.621 |
| | | | | | B.1.596 | AY.34 |
| | | | | | B.6 | AY.54 |

| | | | | | | |
|---|---|---|---|---|---|---|
| | | | | | C.17 | AY.75.1 |
| | | | | | C.35 | AY.80 |
| | | | | | N.4 | AY.92 |
| | | | | | N.5 | B.1.1.306 |
| | | | | | P.1.14 | B.1.177.81 |
| | | | | | P.1.15 | B.1.400 |
| | | | | | P.1.7 | B.1.438.1 |
| | | | | | | B.1.546 |
| | | | | | | B.1.595 |
| | | | | | | B.1.81 |
| | | | | | | P.1.14 |
| | | | | | | A.2.5 |
| | | | | | | AL.1 |
| | | | | | | AY.105 |
| | | | | | | AY.110 |
| | | | | | | AY.116 |
| | | | | | | AY.24 |
| | | | | | | AY.3.1 |
| | | | | | | AY.34.1 |
| | | | | | | AY.35 |
| | | | | | | AY.46.2 |
| | | | | | | AY.46.5 |
| | | | | | | AY.61 |
| | | | | | | AY.69 |
| | | | | | | AY.74 |
| | | | | | | AY.79 |
| | | | | | | AY.91 |
| | | | | | | AY.98.1 |
| | | | | | | B.1.1.141 |
| | | | | | | B.1.1.326 |
| | | | | | | B.1.1.391 |
| | | | | | | B.1.1.416 |
| | | | | | | B.1.1.432 |
| | | | | | | B.1.1.47 |
| | | | | | | B.1.1.521 |
| | | | | | | B.1.147 |
| | | | | | | B.1.177.12 |
| | | | | | | B.1.177.52 |
| | | | | | | B.1.177.60 |
| | | | | | | B.1.199 |
| | | | | | | B.1.236 |
| | | | | | | B.1.355 |

| | | | | | | |
|---|---|---|---|---|---|---|
| | | | | | | B.1.36.16 |
| | | | | | | B.1.36.31 |
| | | | | | | B.1.36.8 |
| | | | | | | B.1.426 |
| | | | | | | B.1.448 |
| | | | | | | B.1.499 |
| | | | | | | B.1.93 |
| | | | | | | C.37 |
| | | | | | | Q.8 |
| | | | | | | A |
| | | | | | | A.1 |
| | | | | | | A.2.2 |
| | | | | | | A.2.3 |
| | | | | | | A.27 |
| | | | | | | A.28 |
| | | | | | | AD.2 |
| | | | | | | AU.3 |
| | | | | | | AY.101 |
| | | | | | | AY.111 |
| | | | | | | AY.117 |
| | | | | | | AY.120.1 |
| | | | | | | AY.121 |
| | | | | | | AY.13 |
| | | | | | | AY.14 |
| | | | | | | AY.16 |
| | | | | | | AY.2 |
| | | | | | | AY.4.4 |
| | | | | | | AY.48 |
| | | | | | | AY.5.3 |
| | | | | | | AY.5.4 |
| | | | | | | AY.59 |
| | | | | | | AY.64 |
| | | | | | | AY.65 |
| | | | | | | AY.67 |
| | | | | | | AY.7 |
| | | | | | | AY.93 |
| | | | | | | B.1.1.125 |
| | | | | | | B.1.1.13 |
| | | | | | | B.1.1.161 |
| | | | | | | B.1.1.192 |
| | | | | | | B.1.1.196 |
| | | | | | | B.1.1.207 |

| | | | | | | |
|---|---|---|---|---|---|---|
| | | | | | | B.1.1.241 |
| | | | | | | B.1.1.273 |
| | | | | | | B.1.1.277 |
| | | | | | | B.1.1.288 |
| | | | | | | B.1.1.317 |
| | | | | | | B.1.1.342 |
| | | | | | | B.1.1.348 |
| | | | | | | B.1.1.359 |
| | | | | | | B.1.1.363 |
| | | | | | | B.1.1.372 |
| | | | | | | B.1.1.389 |
| | | | | | | B.1.1.39 |
| | | | | | | B.1.1.397 |
| | | | | | | B.1.1.40 |
| | | | | | | B.1.110.3 |
| | | | | | | B.1.134 |
| | | | | | | B.1.157 |
| | | | | | | B.1.160.29 |
| | | | | | | B.1.177.15 |
| | | | | | | B.1.177.3 |
| | | | | | | B.1.177.4 |
| | | | | | | B.1.177.41 |
| | | | | | | B.1.177.70 |
| | | | | | | B.1.177.87 |
| | | | | | | B.1.180 |
| | | | | | | B.1.192 |
| | | | | | | B.1.201 |
| | | | | | | B.1.221.2 |
| | | | | | | B.1.258.14 |
| | | | | | | B.1.258.22 |
| | | | | | | B.1.258.3 |
| | | | | | | B.1.260 |
| | | | | | | B.1.265 |
| | | | | | | B.1.270 |
| | | | | | | B.1.298 |
| | | | | | | B.1.3 |
| | | | | | | B.1.311 |
| | | | | | | B.1.325 |
| | | | | | | B.1.328 |
| | | | | | | B.1.336 |
| | | | | | | B.1.337 |
| | | | | | | B.1.36.7 |

| | | | | | | |
|---|---|---|---|---|---|---|
| | | | | | | B.1.36.9 |
| | | | | | | B.1.369 |
| | | | | | | B.1.381 |
| | | | | | | B.1.385 |
| | | | | | | B.1.389 |
| | | | | | | B.1.390 |
| | | | | | | B.1.433 |
| | | | | | | B.1.462 |
| | | | | | | B.1.465 |
| | | | | | | B.1.471 |
| | | | | | | B.1.517 |
| | | | | | | B.1.521 |
| | | | | | | B.1.524 |
| | | | | | | B.1.527 |
| | | | | | | B.1.547 |
| | | | | | | B.1.551 |
| | | | | | | B.1.564.1 |
| | | | | | | B.1.567 |
| | | | | | | B.1.575 |
| | | | | | | B.1.580 |
| | | | | | | B.1.603 |
| | | | | | | B.1.619.1 |
| | | | | | | B.1.628 |
| | | | | | | B.4.8 |
| | | | | | | B.40 |
| | | | | | | B.55 |
| | | | | | | B.6 |
| | | | | | | B.6.6 |
| | | | | | | C.35 |
| | | | | | | C.36.1 |
| | | | | | | N.3 |
| | | | | | | N.4 |
| | | | | | | N.5 |
| | | | | | | P.1.10 |
| | | | | | | P.1.15 |
| | | | | | | P.1.17 |
| | | | | | | P.1.17.1 |
| | | | | | | P.1.7 |
| | | | | | | P.3 |
| | | | | | | Q.1 |
